# Transposable element co-option drives transcription factor neofunctionalization

**DOI:** 10.1101/2025.03.01.640934

**Authors:** Olga Rosspopoff, Danica Milovanović, Sandra Offner, Charlene Raclot, Kelvin Lau, Yoan Duhoo, Julien Duc, Evarist Planet, Claire Damery, Martina Begnis, Florence Pojer, Didier Trono

## Abstract

Gene duplication and domain acquisition are key mechanisms driving protein diversification. However, the impact of these processes on the function and evolution of transcription factors (TFs) remains poorly characterized. Here, we show that the DUF3669 domain, found in a subset of KRAB zinc-finger proteins (KZFPs), originated from the co-option of a fragment of a LINE-1 transposable element ORF1p. Similar to its LINE-1-encoded ancestor, the DUF3669 domain promotes KZFPs trimerization and enables the formation of nuclear condensates with ribonucleoparticle properties, which critically influence the genomic recruitment and action of these TFs. Thus, our study uncovers a direct link between TE co-option and TF neofunctionalization, highlighting how mobile genetic elements shape the evolution of protein functionality.

## Main

Gene duplication is a fundamental driver of genome innovation and adaptation, as first highlighted in Ohno’s seminal work (*1*, *2*). Models such as subfunctionalization, which divides ancestral functions between paralogs, and neofunctionalization, where one copy gains a novel role, describe the evolutionary fate of duplicated genes (*3*, *4*). Functional divergence is critical for the fixation of duplicated genes, typically arising from structural or mutational changes that confer a selective advantage to the host (*5*). Among those, the acquisition of novel protein domains *via* gene fusion, exon shuffling, or domain insertion can rapidly endow duplicated genes with new capabilities (*6–8*). Transposable elements (TEs) play a unique and powerful role in this process. Their intrinsic ability to mobilize and integrate, combined with protein coding potential, positions them as key drivers of host gene evolution (*9*). TE-derived sequences account for roughly half of vertebrate genomes, and their dynamic, species-specific activity has been instrumental in driving evolutionary divergence (*10*, *11*). TEs can notably promote gene diversification by inducing duplication *via* retroposition and by promoting gene fusion through recombination (*5*, *12*). One striking example is the formation of transpo-chimeric gene transcripts, which emerge when TE sequences fuse with host gene exons. These TE-derived fragments are predominantly studied in cases of gene alternative splicing, observed in both physiological and pathological states, including cancer and neurodegeneration (*13–16*). Once viewed solely as selfish elements, TEs are now recognized as assets that can be repurposed to benefit their host, a process coined as TE co-option (*16*).

From an evolutionary standpoint, the contribution of TE-derived fragments to the diversification of host protein-coding genes is likely underestimated. As genes age, evidence of their TE-derived origins is easily obscured by sequence divergence, limiting detection to only a few closely related species over short evolutionary timescales. Nonetheless, TE co-option has been documented across all major eukaryotic lineages, notably in proteins involved in chromatin regulation and transcription factors (TFs) (*17–19*). Yet, the impact of TE co-option in defining the molecular features, functional diversification, and evolutionary trajectories of TFs remains poorly characterized.

To investigate this, we focused on KRAB Zinc Finger Proteins (KZFPs), a vertebrate-specific TF family that exemplifies gene diversification through rapid gene duplication and domain acquisition (*20*, *21*). KZFPs represent the largest family of TFs encoded by mammals, with roughly 400 members in human (*22*, *23*). Their conserved structure features a KRAB domain, which mediates repression via H3K9me3 deposition, alongside a variable zinc-finger (ZF) array that determines DNA-binding specificity (*24*, *25*). Subfunctionalization within KZFPs is evident in the emergence of novel DNA-target specificities driven by amino acid changes in their ZF arrays (*23*). Furthermore, one-third of KZFPs also harbor accessory domains including the DUF3669 (*26*, *27*), found exclusively in a subset of amniote-restricted KZFPs, referred to as DKZFPs (*22*, *28*). Because few protein domains have arisen spontaneously during vertebrate evolution (*29*), we thought that the relatively young DUF3669 offered a rare opportunity to dissect the evolutionary forces shaping its origin, structure, and function. In this study, we trace the emergence of the DUF3669 to a segment of a LINE-1 TE and further demonstrate the functional impact of this co-option event on KZFP structural and molecular features, revealing how TE co-option can lead to TF neofunctionalization.

### The evolutionary persistence of DKZFPs underscores their essential functions

To gain novel insights into the evolutionary trajectory of the DUF3669 and DKZFPs, we conducted a comprehensive screen of 103 vertebrate genomes using hidden Markov models (HMMs) to identify KRAB domains, zinc-finger (ZF) arrays, and DUF3669 motifs (fig. S1A-B, see methods, table S1)(*23*, *28*). DUF3669-associated hits were identified across nearly all vertebrate lineages, except for basal vertebrates such as sea lampreys and most sharks/rays, indicating that DUF3669 predates the emergence of KZFP genes (Fig. 1A). In stark contrast, DKZFPs were exclusively detected in amniotes, appearing in similar numbers in mammals (5 ± 2) and birds (4 ± 1), while reptiles exhibited a broader range (3–14 DKZFPs) driven by lineage-specific expansions in sub-groups such as turtles (fig. S2A). To trace DKZFP evolution, we computed the pairwise identity of their ZF signatures, defined as the set of 4 DNA-contacting residues per ZF within a ZF array (fig. S2B, table S2) (*22*, *23*, *30*). By applying a 60% identity threshold to delineate ortholog groups, we determined that all human DKZFPs originated in mammals except for ZNF777 and ZNF282, which were detected in non-mammalian species and thus represent the oldest traceable KZFP members in amniotes (*22*, *31*). Subtle lineage-specific variations exist within the core set of mammalian DKZFPs. For example, while six *DKZFP* genes were identified in humans, mice, and rabbits, *ZNF783* is specifically absent from the rodent lineage (Fig. 1B). Conversely, *Zfp956* is unique to rodents, exists as a pseudogene in rabbits, and is missing in humans (Fig. 1B). This evolutionary persistence is atypical for KZFPs, observed in less than 5% family members (*27*), which strongly suggests that DKZFPs perform essential, deeply conserved functions. Consistently, four of the six human DKZFPs (ZNF777, ZNF746, ZNF282, and ZNF398) rank among the top 15% of KZFPs most intolerant to loss-of-function variants (Fig. 1C). ZNF777 exhibits a maximal intolerance score of 1, a characteristic often linked to haploinsufficiency and dominant genetic diseases (*32*). Moreover, sequences encoding the N-terminal regions of DKZFPs, including the DUF3669 and KRAB domains, exhibit low frequencies of disruptive mutations such as stop codons or frameshifts (fig. S2C). In summary, DKZFPs are among the most ancient KZFPs detected in amniotes, distinguished by a unique evolutionary trajectory and strong purifying selection against loss-offunction variants in the human population.

**Figure 1.**
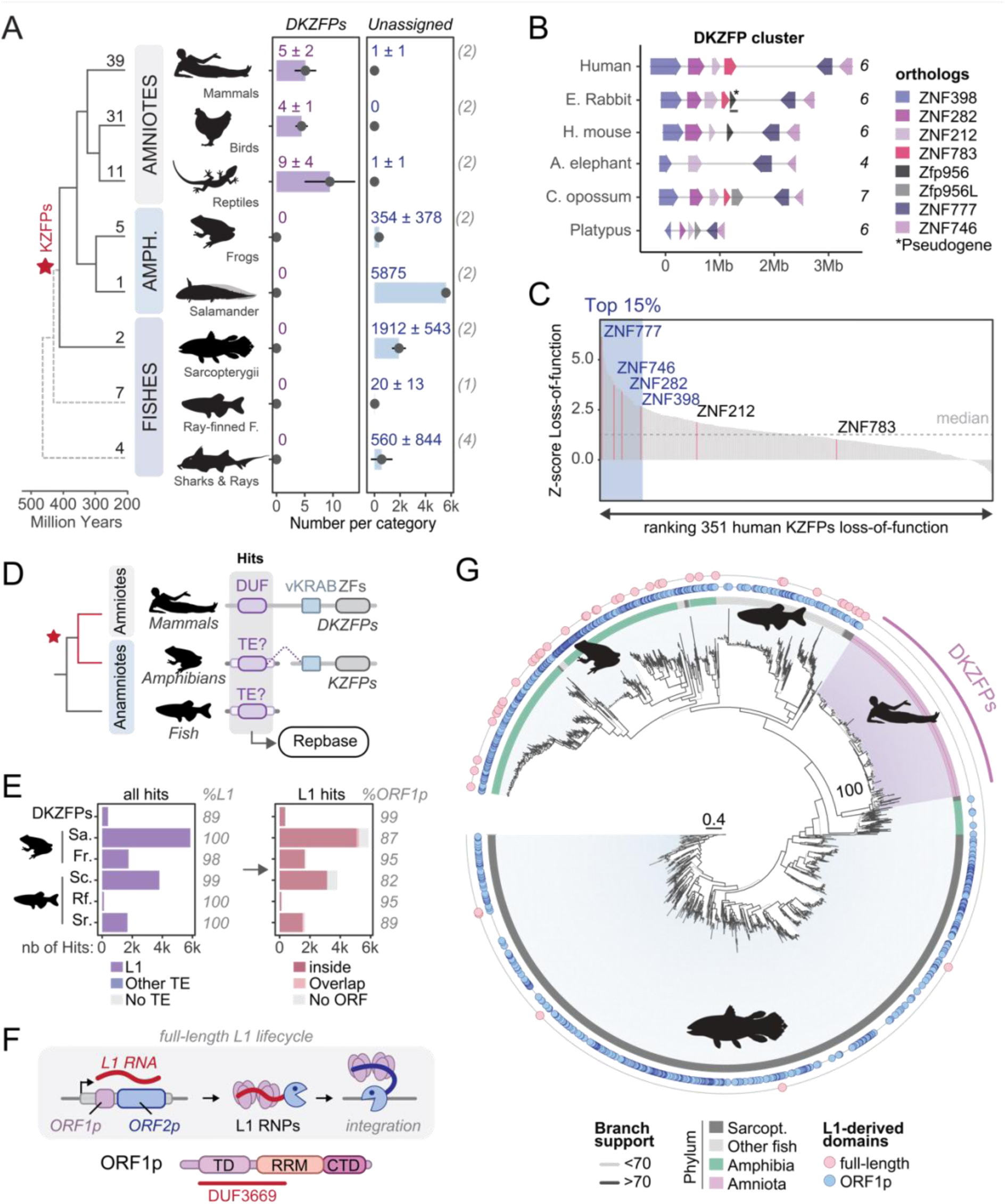
DKZFPs originated from a single co-option event of an ancestral L1 by a *KZFP* gene. A. Evolutionary distribution of DUF3669-associated hits across vertebrate lineages. Italicized numbers in brackets indicate the mean number of ambiguous hits. A red star indicates the emergence of KZFPs. B. Synteny analysis of the DKZFP cluster across selected mammalian species. C. Loss-of-function (LoF) intolerance z-scores of human KZFPs (*32*) with DKZFPs highlighted in pink. The dashed line represents the median LoF z-score for all human genes. D. Schematic representation of DUF3669 annotation using Repbase (*33*). E. Number of DUF3669 hits associated with transposable elements classified by Repbase (left), and the number of L1-associated hits mapped onto the L1-ORF1p consensus (right). Abbreviations: Sa, Salamanders; Fr, Frogs; Sc, Sarcopterygii; Rf, ray-finned fish; Sr, sharks and rays. F. Schematic representation of L1 retrotransposition (top) with structure of the L1-ORF1p domain most frequently associated with the DUF3669 domain. Abbreviations : RNPs, ribonucleoparticles; TD, trimerization domain; RRM, RNA-recognition Motif; CTD, C-terminal domain. G. Maximum likelihood phylogenetic tree of DUF3669-associated hits.

### DUF3669 emerged from the co-option of a LINE-1 element

Beyond their association with DKZFPs, DUF3669 hits showed an unexpectedly high number of occurrences in anamniote species, notably salamanders (axolotl; n = 5’875, Fig. 1A), with most located in intergenic regions (88.5%, fig. S3A). To test if novel DUF3669 hits emerged *via* tandem duplication, a known driver of *KZFP* gene expansion (*20*, *23*), we analyzed their linear genomic proximity (fig. S3B, see methods). In contrast to DKZFPs, most anamniote hits appeared as isolated entities; only 23% of axolotl hits formed clusters averaging 2.28 hits per cluster. Since these DUF3669 hits were largely non-genic and interspersed, we examined if they showed sequence similarity to TEs using Repbase (*33*) (Fig. 1D, table S1). DUF3669 hits were unequivocally matched to LINE-1 (L1) elements (89-100%, Fig. 1E), a class of TEs that self-replicate *via* a “copy and paste” mechanism using an RNA intermediate (*34*) (Fig. 1F). L1s have significantly diversified in vertebrates, with major structural changes within their protein domains at the amniote-anamniote split (*35*). Indeed, 93.97% of L1-annotated DUF3669 hits matched L1 subfamilies specific to anamniotes, but not to mammals or amniotes (fig. S3C).

To identify which L1 protein domains were incorporated in DKZFPs, we mapped DUF3669 hits onto consensus sequences of L1 subfamilies. Full-length L1s encode two proteins: ORF1p and ORF2p (Fig. 1F). ORF2p has endonuclease and reverse transcriptase activities, whereas ORF1p acts as an RNAbinding chaperone that typically contains a trimerization domain (TD), an RNA-recognition motif (RRM), and a C-terminal domain (CTD). Most DUF3669 hits mapped to the L1-ORF1p consensus (82– 99%, Fig. 1E), with the greatest overlap involving the TD and adjacent RRM-coding sequences (fig. S3D). Notably, the highest conservation was found in a 19–amino acid segment overlapping the TD-RRM junction (entropy score of 0.63, fig. S3D). This prompted us to investigate the presence of L1-derived domains within the ±10 kb of DUF3669 hits. L1-derived elements were enriched in the vicinity of DUF3669 hits in anamniotes but not in that of DKZFPs (fig. S3E). Finally, a maximum-likelihood phylogenetic tree revealed that DKZFP-associated hits formed a single monophyletic group, indicating a single co-option event of an L1 into a KZFP gene (branch support 100%, Fig. 1G). The short branch lengths associated with DKZFPs suggests the cessation of a prototypical pattern of L1 diversification, consistent with the fixation of an L1derived domain as a functional component in the host genome. In summary, our findings reveal that DUF3669 emerged from an ancient L1 element that flourished 320 million years ago in the common ancestor of amniotes and amphibians. In amniotes, this TE-derived domain was exapted into a KZFP gene and fixed under strong purifying selection, while in anamniotes it continued to propagate as an active L1.

### L1-derived domain co-option enabled KZFP trimerization

We hypothesized that the co-option of an L1 element endowed KZFPs with novel molecular properties. To explore structural homologies between L1-ORF1p and host-encoded DKZFPs, we investigated whether the DUF3669 domain could promote trimeric assembly, a supramolecular structure rarely observed among TFs (*36*, *37*). We first examined the trimerization properties of a purified recombinant fragment containing only the DUF3669 and KRAB domains (DK; residues 1–191) of ZNF746, also known as PARIS (Fig. 2A and fig. S4A). Analysis by native mass spectrometry, mass photometry, and size-exclusion chromatography with multi-angle light scattering (SEC-MALS) confirmed that the DK fragment forms an exclusive trimer *in vitro* (~66 kDa), with no detectable dimeric or monomeric forms. Consistent with a trimeric coiled-coil conformation, circular dichroism spectroscopy of the DK fragment revealed a prototypical α-helical spectrum (ellipticity ratio of 0.98; fig. S4D). Sequence analysis further supported a coiled-coil structure by identifying 10 consecutive canonical heptads, along with two RhxxhE trimerization motifs spanning heptads I and VIII (fig. S3D–S4E), features that are also shared with anamniote L1s (*35*). To further validate our findings, we performed cryogenic electron microscopy (Cryo-EM) on the recombinant DK trimer. Despite the challenges posed by its small size, unsupervised alignment of 2D classes successfully generated a low-resolution surface map (Fig. 2B-C and fig. S5A, see methods), representing a significant advancement in structural analysis for complexes of this scale. The DK trimer adopts an extended rod-like conformation with an overall length of 142 Å, with all three KRAB domains located on one side of the bundle, with minimal deviations from the AlphaFold-Multimer (*38*) simulation (R = 0.81; fig. S5B). The triad of KRAB domains formed few interchain contacts and exhibited three-fold rotational symmetry around the supercoil axis (diameter ~64.5 Å; Fig. 3D–E), reminiscent of the RRM/CTD positioning within human L1-ORF1p (*39*, *40*). A second *ab initio* surface map confirmed the presence of three chains within the rod-like structure but lacked features associated to the KRAB positioning (fig. S5D). Two hydrophobic patches were identified within the coiled-coil (<3.5 Å): the C-terminal Contacting Region (CCR) and the Inner Contacting Region (ICR; fig. S5E). Hydrophobic to hydrophilic substitutions within these regions destabilized the trimerization of both the DK fragment and full-length ZNF746 in cells, resulting in dimers and monomers (CCR_MUT_, ICR_MUT_ and combined, Fig. 2F). Notably, ICR mutations had the greatest destabilizing effect, corresponding to positions with minimal sequence variation across DKZFPs (fig. S5F). Thus, functional diversification of DKZFPs results from amino acid heterogeneity within the CCR, as well as the KRAB domain and unique ZF arrays (fig. S5F). Regardless of these sequence variations, pairwise co-immunoprecipitation revealed that all human DKZFPs are capable of homo- and hetero-oligomerization (Fig. 2G) in a DUF3669-dependent manner (fig. S5G)(*27*, *28*). Therefore, *in vivo* association of DKZFPs is likely driven by their relative abundance within cells, influencing trimer composition and stoichiometry in a tissue-specific manner, as highlighted by their unique expression patterns across of human fetal tissues (*41*) (Fig. 2H). Taken together, our results demonstrate that the cooption of a L1 segment endowed KZFPs with the capacity to form both homo- and hetero-trimers, thereby opening the way to combinatorial regulatory activities.

**Figure 2.**
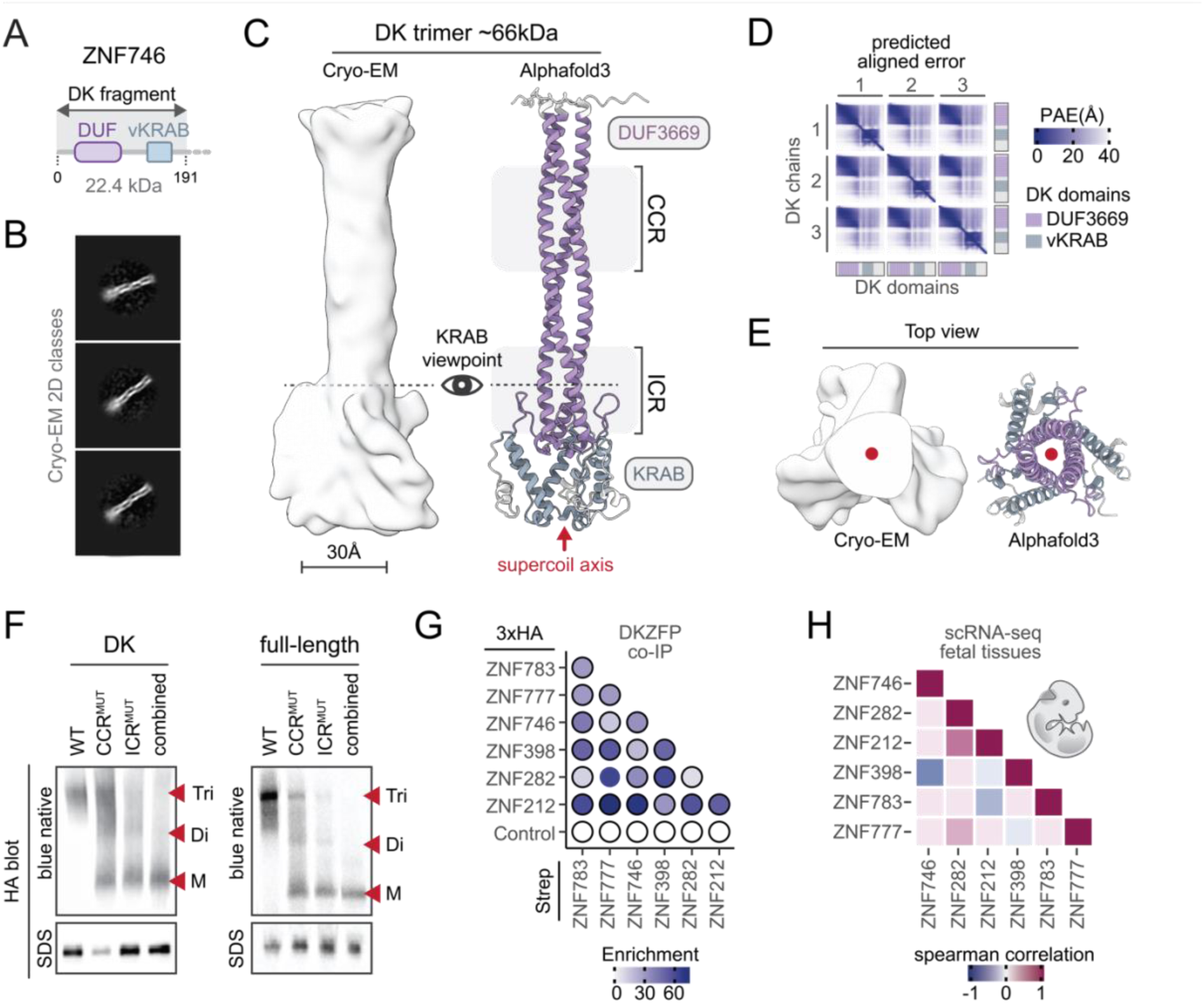
L1 co-option triggered homo- and hetero-trimerization of KZFPs. A. Schematic representation of the recombinant DK fragment of ZNF746 produced in E. coli. B. Representative pictures of Cryo-EM 2D classes of ZNF746 DK fragment. C. Cryo-EM density map (left) and AlphaFold3-Multimer model (right) of the DK fragment. CCR, C-terminal Contacting Region; ICR, Inner Contacting Region. D. Predicted distances of the Aligned Error of the AlphaFold3-Multimer model across the three DK chains, marking the DUF3669 and KRAB domain positions. E. Top view of the KRAB domains from the supercoil axis at the KRAB viewpoint (red dot). F. Western blot with anti-HA antibody on Blue native (top) and SDS-PAGE (bottom) gels of whole-cell extracts from HEK293T cells transfected with HA-tagged DK fragment (left) or full-length ZNF746 (right), either wild-type (WT) or with CCR, ICR, or combined mutations. Tri: trimer; Di: dimer; M: monomer. G. Dot plot quantifying coimmunoprecipitation assays with HA- and Strep-tagged constructs of six human DKZFPs. LacZ-3xHA and GFPStrep serve as controls. H. Heatmap showing Spearman correlation of human DKZFP expression levels across 16 fetal tissues (*41*)

**Figure 3.**
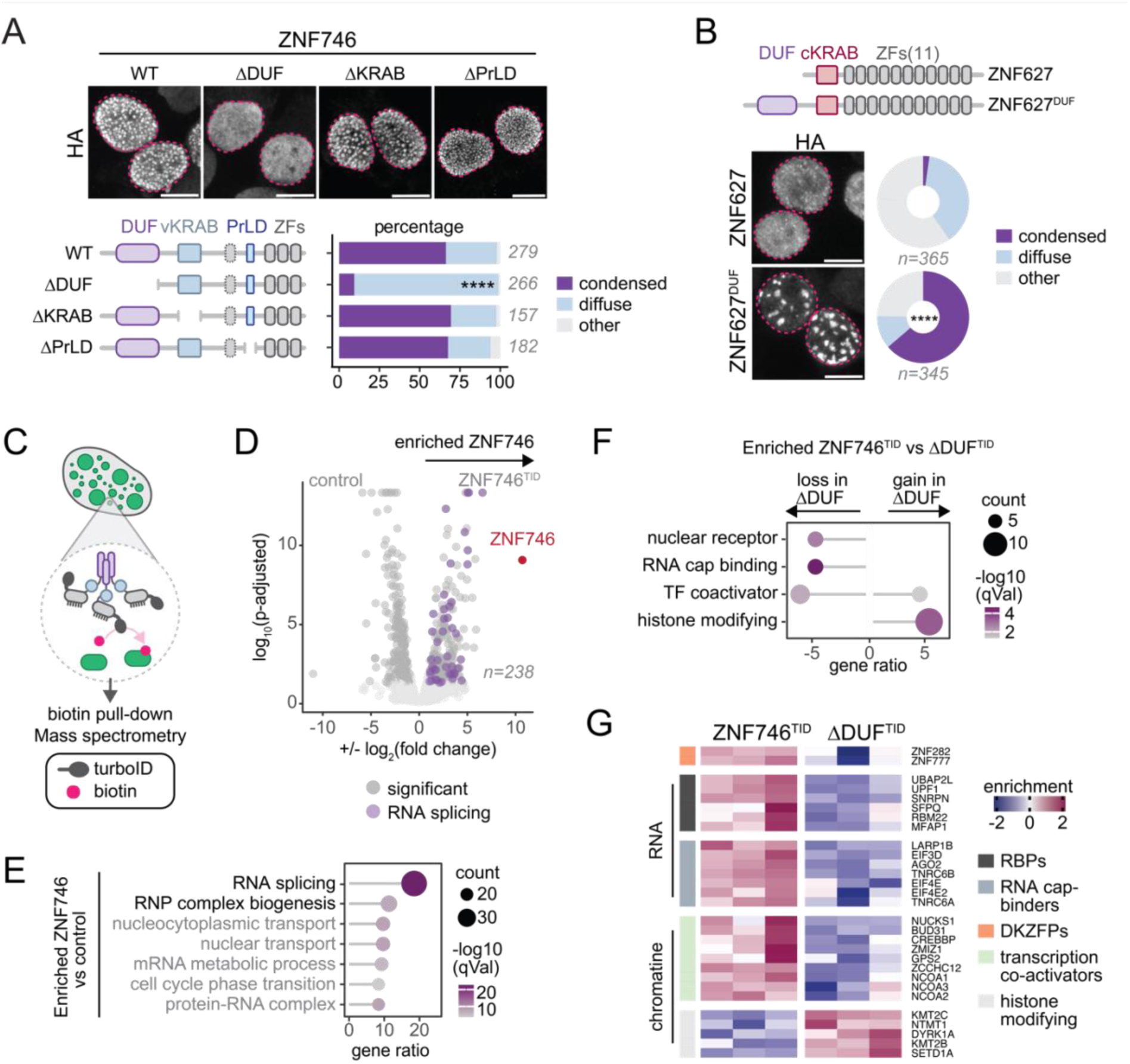
L1 co-option allowed DKZFP to form condensate, rich in RBPs and transcriptional coactivators. A. Representative immunofluorescence images (z-projection) of HA-tagged ZNF746 and associated mutants (top) with quantification of condensate formation (bottom). The pink dotted line delineates the nucleus as assessed by DAPI staining. Numbers above the bar plot indicate the number of HEK293T cells counted (chi-square test). Scale bar : 10μm. B. Same as A with HA-tagged ZNF627 and ZNF627 fused with a C-terminal DUF3669 domain. C. Scheme of proximity biotinylation for full-length (ZNF746^TID^) and DUF-deleted ZNF746 (ΔDUF^TID^) fused to TurboID. D. Volcano plot showing log₂ fold changes between ZNF746^TID^ and mock-treated HEK293T cells with respect to log₁₀ adjusted p-values. The gray dotted lines indicate thresholds (fold change of 1.5 and p-adjusted 0.05). E. Enriched gene ontologies for ZNF746^TID^ compared to mock-treated control cells. F. Enriched gene ontologies for ZNF746^TID^ compared to ΔDUF^TID^. G. Heatmap of top proteins significantly depleted in ΔDUF^TID^ compared to ZNF746^TID^.

### DUF3669 drives the formation of condensates analogous to ribonucleoparticle

As ZNF746 forms DUF3669-mediated trimers, we investigated whether this property translates into distinct subcellular localization patterns. In human L1, ORF1p trimerization coupled with nucleic acid binding drives the assembly of ribonucleoprotein particles (RNPs) into biomolecular condensates essential for retrotransposition (Fig. 1F)(*42*, *43*). We found overexpression of HA-tagged ZNF746 to induce nuclear condensate formation, a pattern lost upon deletion of the DUF3669 domain, consistent with it requiring trimerization (Fig. 3A and fig. S6A, See supplementary text). In contrast, both the KRAB domain and the Prion-Like Domain (PrLD), a low-complexity region implicated in ZNF746 aggregation (*44*, *45*), were dispensable for initial condensation and subcellular localization (Fig. 3A). Noteworthy, all human DKZFPs could form higher-order assemblies in a DUF-dependent manner, albeit along different patterns (fig. S6B). Further supporting the essential role of DUF3669 in DKZFP biology, when it was appended to ZNF627 (ZNF627^DUF^), a KZFP sharing little sequence homology with DKZFPs (*23*), it shifted the nuclear distribution of this protein from diffuse to condensed (Fig. 3B and fig. S6C).

Condensates locally concentrate proteins, DNA, and RNA to coordinate diverse molecular processes (*46*, *47*). To examine the composition of DUF3669-mediated condensates, we performed a proximity biotinylation assay by fusing DKZFPs to the biotin ligase TurboID (Fig. 3C, table S3) (*48*). We first observed that fusing TurboID to full-length ZNF746 (ZNF746^TID^) or a DUF-deleted variant (ΔDUF^TID^) did not alter the subcellular localization or condensation patterns of these proteins (fig. S6D). Proteomic analysis revealed that proteins significantly enriched in ZNF746 samples versus mock-treated control were primarily involved in mRNA splicing (p-adjusted = 1.9E-22) and RNP complex biogenesis (p-adjusted = 1.7E-8, Fig. 3D-E), indicating that ZNF746 condensates are rich in RNA-binding proteins (RBPs). Deletion of the DUF3669 resulted in a marked depletion of endogenous DKZFPs in the proximity of ZNF746, including ZNF282 and, to a lesser extent, ZNF777, reflecting the relative abundance of these proteins in the cell line used for this experiment (fig. S6E-F). Additionally, RNA processing factors (Fig. 3F-G), including the RNA helicase UPF1, a well-known L1-ORF1p interactor (*49*, *50*), were lost from the environment of ZNF746 upon DUF3669 deletion. DUF3669-dependent proximal proteins also included several transcriptional coactivators involved in developmental signaling pathways (e.g., TGF-β signaling and nuclear receptors) (*51*), and components of histone-modifying complexes (Fig. 3F-G, table S4). Furthermore, condensate composition varied among DKZFPs; for example, ZNF282 displayed both overlapping and unique DUF3669-dependent proximal proteins (fig. S6G–S6H, table S5), underscoring functional diversification within the family. In sum, these data reveal that the DUF3669 acts as a molecular scaffold, partitioning ZNF746 function into RNA-rich condensates, analogous to L1-ORF1p RNPs, that are further specialized in chromatin-related processes.

### The DUF3669 domain expands ZNF746 chromatin association

To assess the functional impact of L1-derived neofunctionalization on a core attribute of DKZFPs, we examined ZNF746 chromatin occupancy *in situ*. Profiling of HA-tagged ZNF746 binding by Cut&Tag (*52*) revealed a marked enrichment at promoter regions (~44%, p-value < 0.001, Fig. 4A) and a strong enrichment with histone marks linked to active transcription, such as H3K27Ac and H3K4me3 (fig. S7A). In contrast to most other KZFPs, ZNF746 recruitment sites showed no enrichment for the repressive H3K9me3 mark (*22*), suggesting that neofunctionalization entails a shift from canonical repression to association with transcriptionally active regions. Consistently, H3K4me3 foci colocalized at the periphery of ZNF746 condensates (Fig. 4B), hinting that these condensates physically cluster active genomic loci. Furthermore, ZNF746 binding sites display a range of overlap with other DKZFPs (fig. S6E). Approximately one-third of ZNF746 binding sites overlapped with those of ZNF282 (31.5%, fig. S7B), whereas cooccurrence with ZNF777 was minimal (fig. S7C). These results suggest that ZNF746 predominantly engages chromatin as homotrimers rather than as heterotrimeric complexes with other DKZFPs.

**Figure 4.**
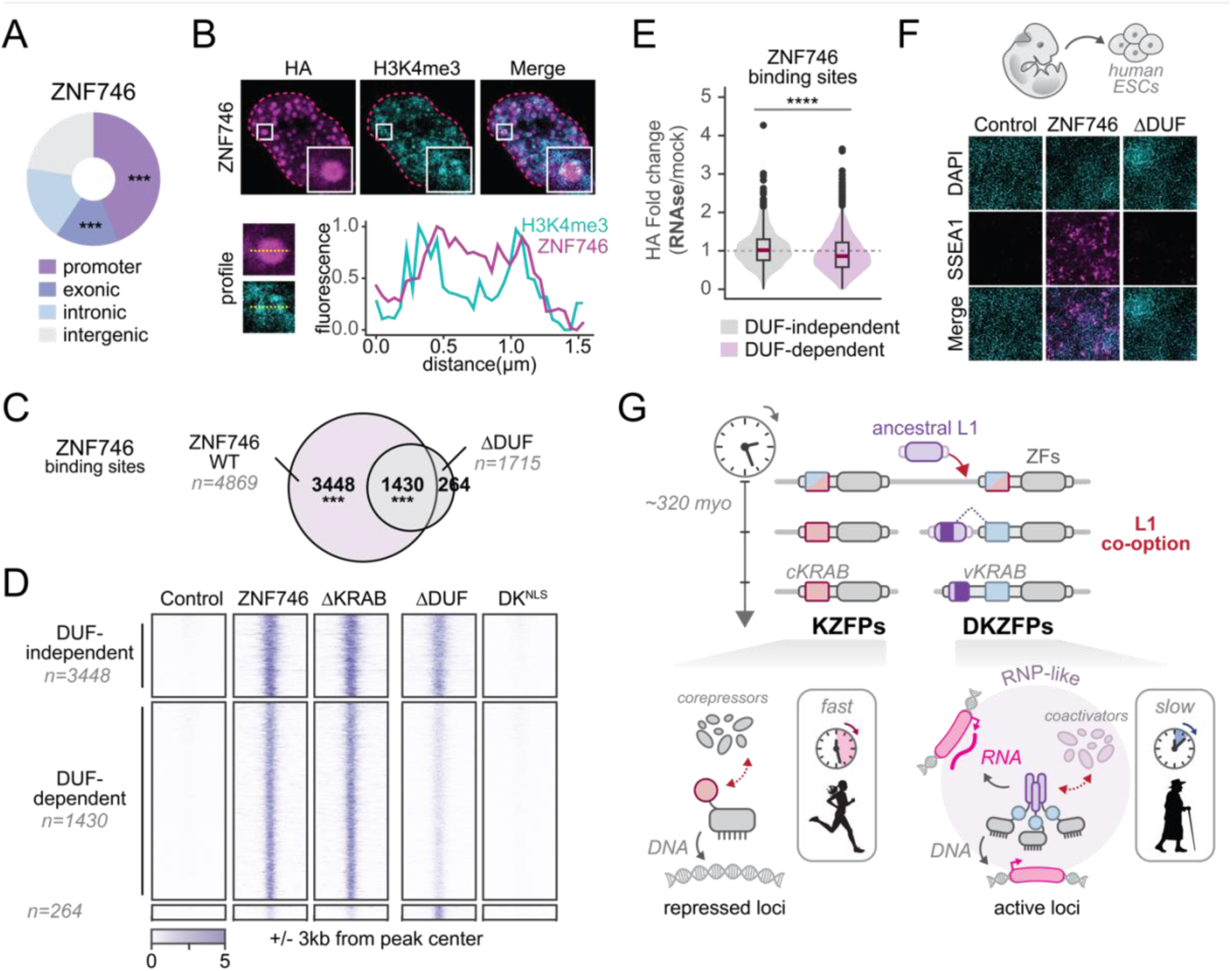
The DUF3669 dictates DKZFP genomic affinity and functionality. A. Pie chart showing the distribution of ZNF746 binding sites across genomic features (permutation test). B. (Top) Representative images and (bottom) profile analysis of ZNF746 colocalization with H3K4me3 in HEK293T cells. The pink dotted line delineates the nucleus, as assessed by DAPI staining. C. Euler plot depicting the overlap of Cut&Tag peaks for HA-tagged ZNF746 and ZNF746ΔDUF (permutation test). D. Heatmap of Cut&Tag signals for ZNF746 and its associated mutants across the indicated lists of ZNF746 peaks. E. Violin plot showing the fold change in HA Cut&Tag signal in control versus RNase A-treated samples for the indicated lists of ZNF746 peaks (Wilcoxon rank-sum test). F. Representative images of SSEA-1 staining after 7 days of overexpression of the indicated constructs in human embryonic stem cells. G. Proposed model of DKZFP neofunctionalization following L1 co-option.

Next, we interrogated whether the DUF3669 influences ZNF746 binding landscape. Strikingly, deletion of DUF3669 reduced by nearly three-fold ZNF746 chromatin association (4869 vs. 1715 peaks, Fig. 4C-D), yet this domain was insufficient in itself to promote chromatin occupancy, as evidenced by the trimeric DK^NLS^ fragment (DK^NLS^, Fig. 4C). In contrast to the DUF3669 deletion, deleting the KRAB domain of ZNF746 did not globally affect its genomic recruitment (ΔKRAB, Fig. 4C and fig. S7D). Therefore, the DUF3669 modulates ZNF746 chromatin affinity with active chromatin regions rather than directly mediating DNA binding, which mostly stems from the zinc finger array. As ZNF746 condensates are enriched in RBPs, we hypothesized that RNA might facilitate chromatin association at DUF3669-dependent sites (*53*). Supporting this model, Cut&Tag with and without prior RNase-A treatment (*54*) revealed that ZNF746 chromatin association was reduced after RNA digestion, with a stronger effect at DUF3669-dependent binding sites (Fig. 4E and fig. S7E). Thus, RNA contributes to ZNF746 chromatin affinity within RNP-like condensates.

Finally, we sought to address whether the DUF3669 affects ZNF746 functionality and its role in transcriptional regulation. First, luciferase assays revealed that the DUF3669 confers activating properties not only to ZNF746 but also to ZNF627(fig. S7F), a *bona fide* repressive KZFP (Fig. 3B) (*23*). This was consistent with the DUF3669-mediated proximity to transcriptional coactivators, observed for ZNF746 and ZNF282 (Fig. 3G). Second, the analysis of genes proximal to ZNF746 binding sites showed a marked enrichment near targets responsive to key signaling pathways, notably TGF-β and WNT (fig. S7G). To substantiate these observations, we turned to human embryonic stem cells (hESCs), a system in which signaling pathways tightly regulate the balance between pluripotency and differentiation. Overexpression of ZNF746 triggered ectopic differentiation of hESCs (SSEA-1 positive cells), an effect strictly dependent on the DUF3669 (Fig. 4F). Furthermore, treatment of ZNF746-overexpressing cells with a TGF-β inhibitor effectively abrogated cell differentiation (fig. S7H), consistent with a direct involvement of the DUF3669 in mediating ZNF746 transcriptional regulation downstream of TGF-β signaling. Collectively, these data demonstrate that the co-option of an L1-ORF1p segment endows KZFPs with enhanced chromatin affinity that critically influences their function as transcriptional activators, highlighting the transformative impact of TE exaptation on TF evolution through neofunctionalization.

## Discussion

Owing their young evolutionary age, rapid expansion and extensive target diversification, KZFPs offer a unique setting to investigate the fate of duplicated TF genes in real time. Although neofunctionalization is a compelling and well-supported concept, its overall prevalence remains a matter of debate. Similarly, the biological relevance of TE-host gene fusion is often questioned: some events are confined to a few closely related species, while others are so ancient that identifying the functional shifts resulting from the exaptation of TE components becomes challenging. Our data provide, with unprecedented detail, evidence that the capture of a segment of L1-ORF1p by a KZFP gene gave rise to DKZFP derivatives endowed with a unique structural scaffold, activating functions, and modes of genomic recruitment that distinguish them from the majority of KZFP family members. Thus, this co-option event, which occurred at the dawn of amniotes approximately 320 million years ago, demonstrates that the acquisition of a TE-derived domain can offer a fast-track route to genetic innovation among duplicated genes (Fig. 4G). DKZFPs stand out among KZFPs by exhibiting an evolutionary trajectory characterized by slow turnover and deep conservation across mammals, underscoring their essential roles in the biology of these species, including human. They possess structural features that promote both homo- and hetero-trimerization, leading to the formation of higher-order assemblies, and they employ novel modes of genomic recruitment that diverge from classical zinc finger-mediated, sequence-specific DNA recognition. Moreover, they associate with transcriptional coactivators rather than repressors, a distinction supported by a KRAB domain that lacks traditional silencing function (Fig. 4H) (*27*, *55*). The coexistence and evolutionary persistence of both ancestral (KZFP) and novel (DKZFP) functions in extant genomes support the concept of TE-driven neofunctionalization of TFs (*4*). DKZFPs continued to diversify after their initial emergence, as evidenced by differences in their genomic targets and associated functional effectors, thereby establishing a subfamily of proteins that are predicted to act in combinatorial fashions to produce a range of functional outcome. Beyond TFs, our findings suggest that TE co-option events may have reshaped the diversification of L1 elements throughout vertebrate evolution (*35*). In amniotes, DUF3669 hits are exclusively linked to DKZFP genes, whereas in anamniotes they remain associated with L1-derived sequences. This pattern implies a genetic incompatibility between DKZFPs and the ancestral L1 elements from which the DUF3669 domain originated, with coexistence being detrimental to either the host or the retroelement itself. Finally, L1 elements account for about 20% of the human genome and remain among the few retrotransposons capable of autonomous mobilization in our species (*34*). L1-derived sequences are frequently incorporated into cellular RNAs (*56*), thus representing of latent reservoir of functional protein domains (*57*) and proteins (*58*). The functional implications of L1 co-option into the pool of protein-coding genes of the host may thus extend far beyond DKZFPs.

## Acknowledgments

We thank past and present members of the laboratory of Virology and Genetics, particularly Romain Forey, Olimpia Bompadre and Arianna Dorschel for their valuable input on the manuscript and their feedback throughout the years. We are grateful to Prof. Aleksandar Antanasijević for his assistance in refining the cryo-EM structure and to Prof. Cedric Feschotte for his insightful comments.

We also acknowledge the EPFL Gene Expression Core Facility and the Bio Imaging Optics Core Facility for providing access to sequencing and imaging resources. Finally, we thank Natalia Gasilova for supplying data from native-like mass spectrometry.

## Author contributions

Conceptualization, methodology and visualization: O.R. Formal analysis and Data Curation: O.R. and J.D. Investigation: O.R., D.M., S.O., Y.D., K.L. and M.B. Supervision: O.R. and D.T. Funding acquisition: D.T. Writing – Original Draft: O.R., D.M. and D.T. Writing – Review & Editing: O.R., D.M., S.O., Y.D., K.L., F.P., M.B. and D.T.

---

**Figure S1.**
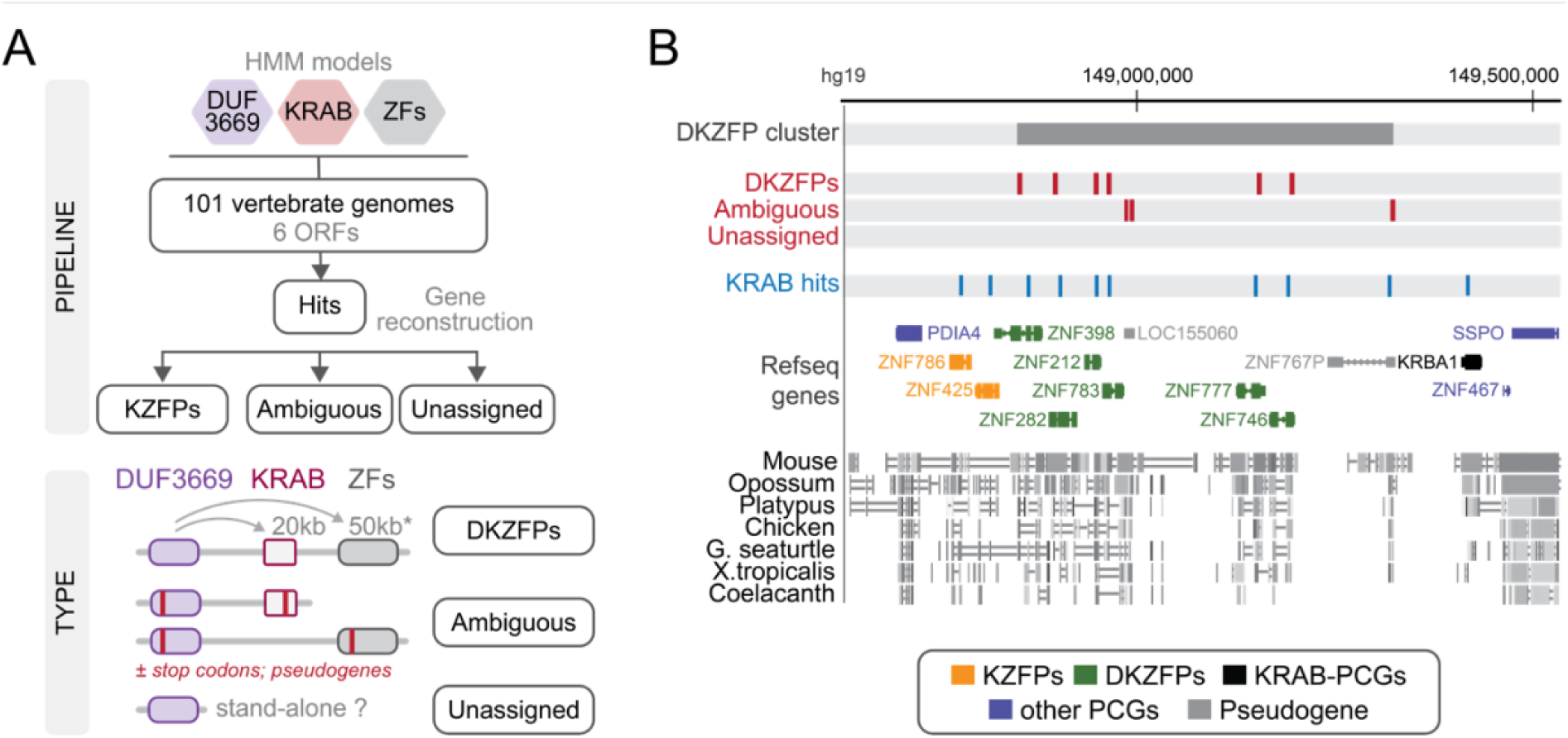
Annotation of DKZFPs in vertebrate genomes. Workflow for classifying DUF3669-associated hits across vertebrates. Hits are classified as DKZFPs, ambiguous, or unassigned based on the arrangement of DUF3669, KRAB, and ZF domains. B. Organization of the human DKZFP gene cluster on chromosome 7, showing the positions of hits corresponding to the DUF3669 and KRAB domains. The bottom track indicates the chained alignments across vertebrate species. Abbreviations: PCGs, protein-coding genes.

**Figure S2.**
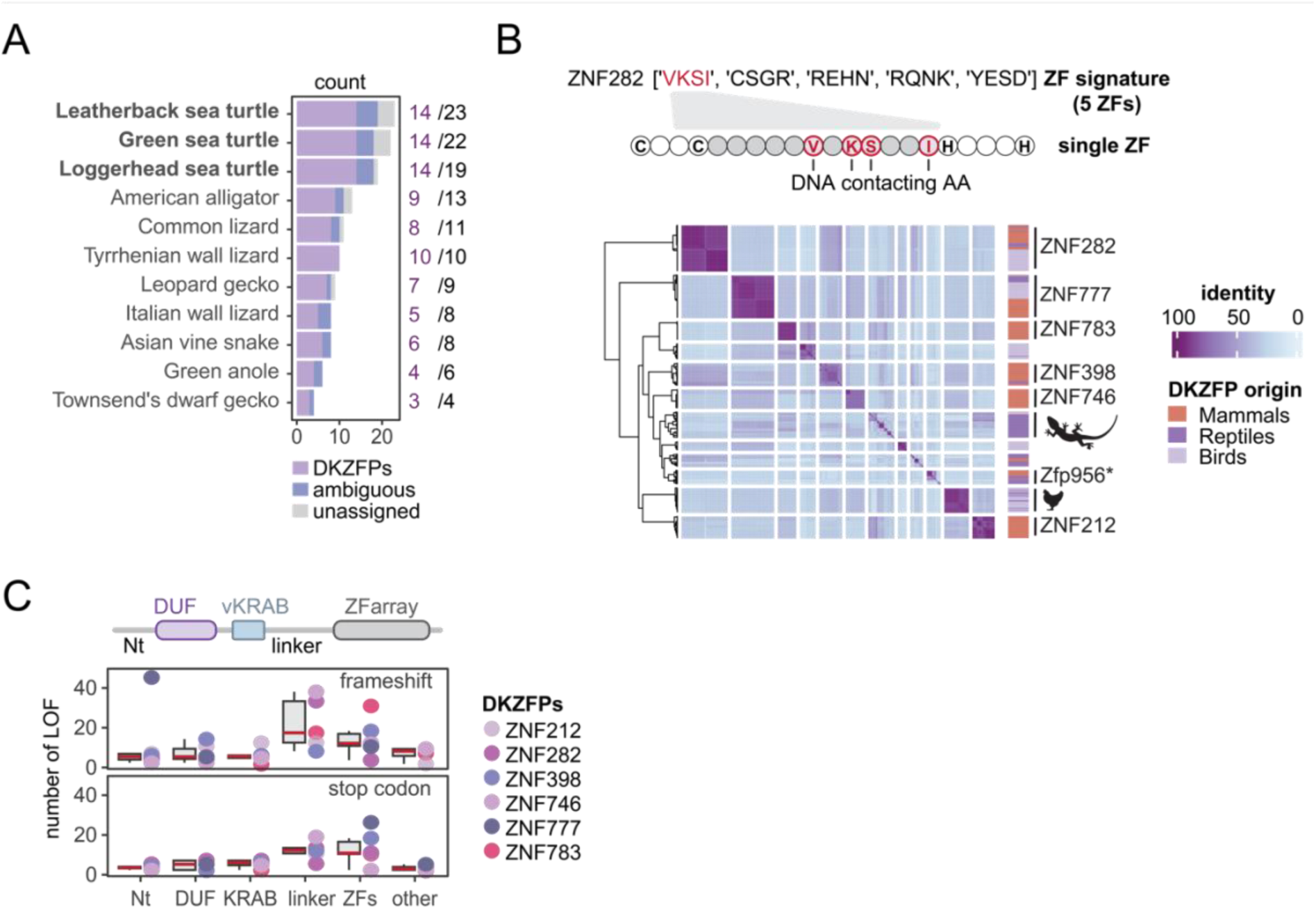
Evolution of DKZFPs in vertebrate genomes. Species-specific expansion of DUF3669 hits across reptile genomes. Numbers in purple indicate the number of DKZFPs, while numbers in black represent the total number of hits. B. Schematic representation of a C2H2 ZF domain structure and its DNA-contacting amino acid residues. The heatmap shows pairwise sequence identity scores of these residues across DKZFPs from different species. Rows and columns represent individual DKZFPs, clustered by identity. C. Box plots showing the percentage of loss-of-function (LoF) variants across distinct DKZFP domains and proteins.

**Figure S3.**
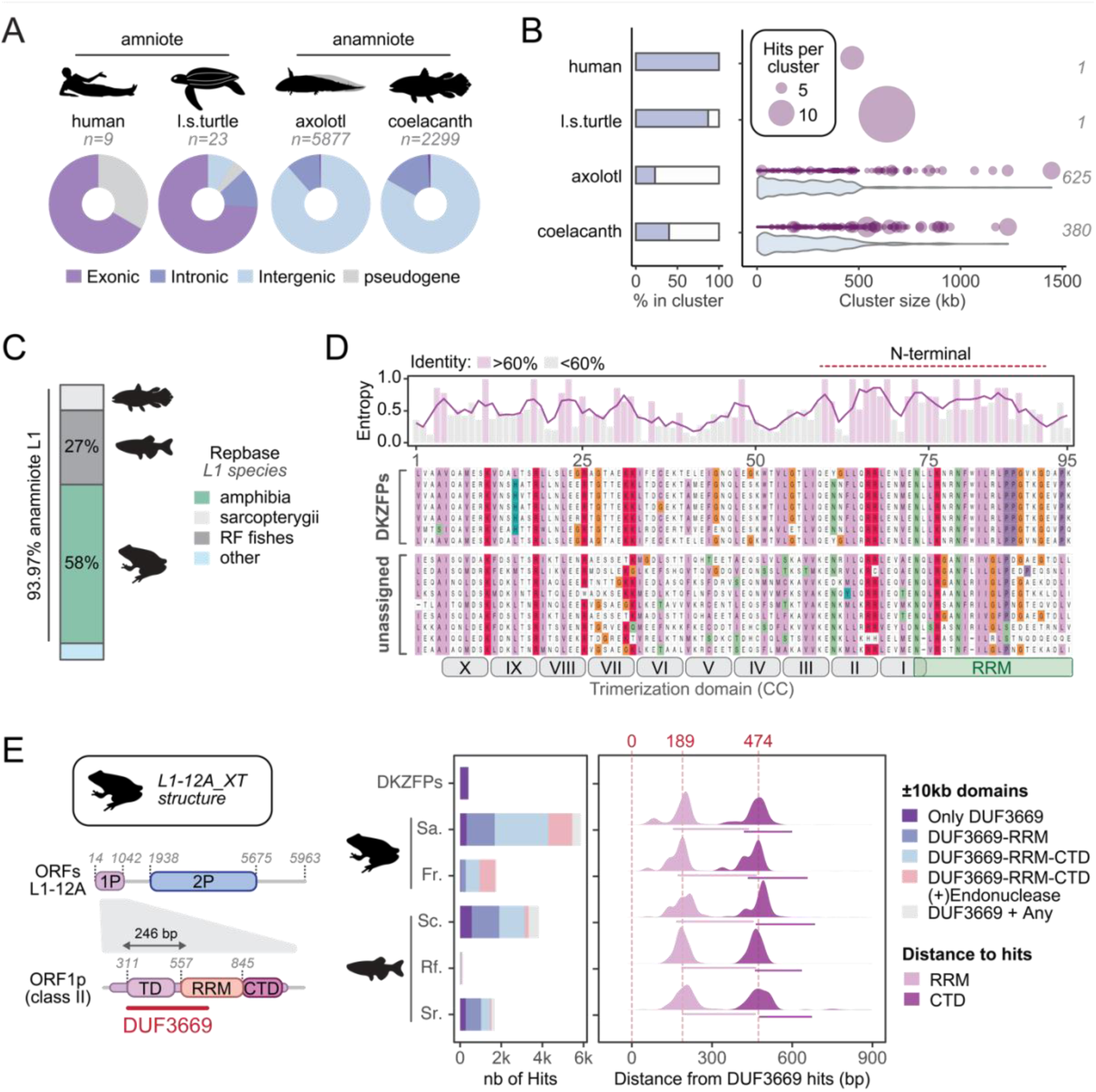
DUF3669 hits are abundant and interspersed elements in anamniotes corresponding to LINE-1. A. Overlap of DUF3669 hits with genomic features in the indicated species. n represents the total number of hits. B. Genomic clustering of DUF3669 hits. The bar chart (left) shows the percentage of hits in clusters (≥2 hits within 500 kb), while the dot plot (right) displays the number of hits per cluster (dot size) and the cluster size in kilobases (x-axis). Italicized numbers indicate the number of clusters identified in each species. C. Bar plot showing the species classifications of L1-associated hits from Repbase. D. Multiple sequence alignment (MSA) of selected DUF3669 hits showing the position of the L1-derived domain. The bar chart above indicates relative Shannon entropy scores, with consensus sequence letters shown where identity exceeds 60%. E. Schematic representation of L1-12A from Xenopus Tropicalis (left) and bar plot (right) depicting the number of L1-associated protein domains within ±10 kb of the DUF3669 hits. The density map displays ORF1p-associated domain start positions relative to DUF3669 hits. The red dotted line indicates the mean distance to the RRM and CTD domains. Abbreviations: Sa, Salamanders; Fr, Frogs; Sc, Sarcopterygii; Rf, ray-finned fish; Sr, sharks and rays.

**Figure S4.**
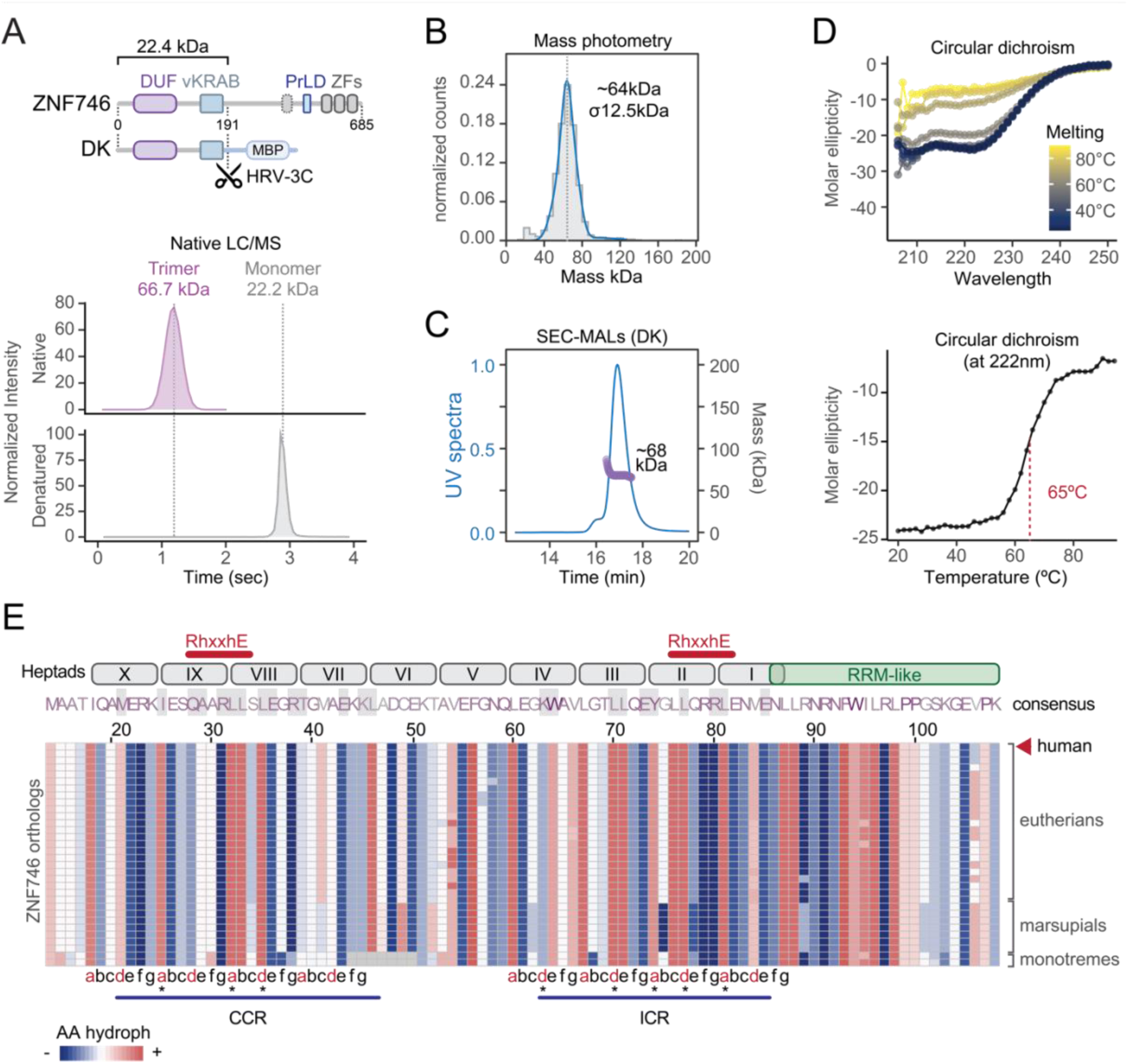
Characterization of ZNF746 DK trimers. A. Schematic representation of the ZNF746 protein domains, including the DUF3669, KRAB, PrLD (Prion-like domain) and ZF array, along with the DK fragment fused to Maltose Binding Protein (MBP) used for purification. Cleavage of the MBP-tag is made by a HRV-3C protease, as indicated by the scissors icon. Density plots at the bottom illustrates LC-MS spectra obtained for the native and denatured DK fragment, showing distinct peaks for the trimeric (native) and monomeric (denatured) forms. B. Mass photometry of the DK fragment. σ represents the standard deviation. C. Size exclusion chromatography coupled with Multi-angle light scattering (SEC-MALS) of the DK fragment. The plot shows molar mass measurements (purple dots, kDa) corresponding to each SEC peak (UV trace in blue). D. Circular dichroism (CD) spectra of the DK fragment at various temperatures, highlighting typical α-helical coiledcoil characteristics with double minima at 209 and 222 nm (top). Melting curve displaying molar ellipticity at 222 nm at different temperatures, indicating a single transition step associated with denaturation. E. Multiple sequence alignment (MSA) of ZNF746 orthologs in mammals. The amino acid sequence corresponds to ZNF746 consensus sequence among orthologs. Grey-shaded areas represent contacting amino acids within 3,5 Å. Heptad numbering (I to X) is shown at the top, with critical RhxxhE trimerization motifs marked by thick red lines. The ICR (Inner Contacting Region) and CCR (C-terminal Contacting Region) heptads are labeled, with asterisks denoting a and d amino acids mutated into glutamine. Related to Fig. 2F.

**Figure S5.**
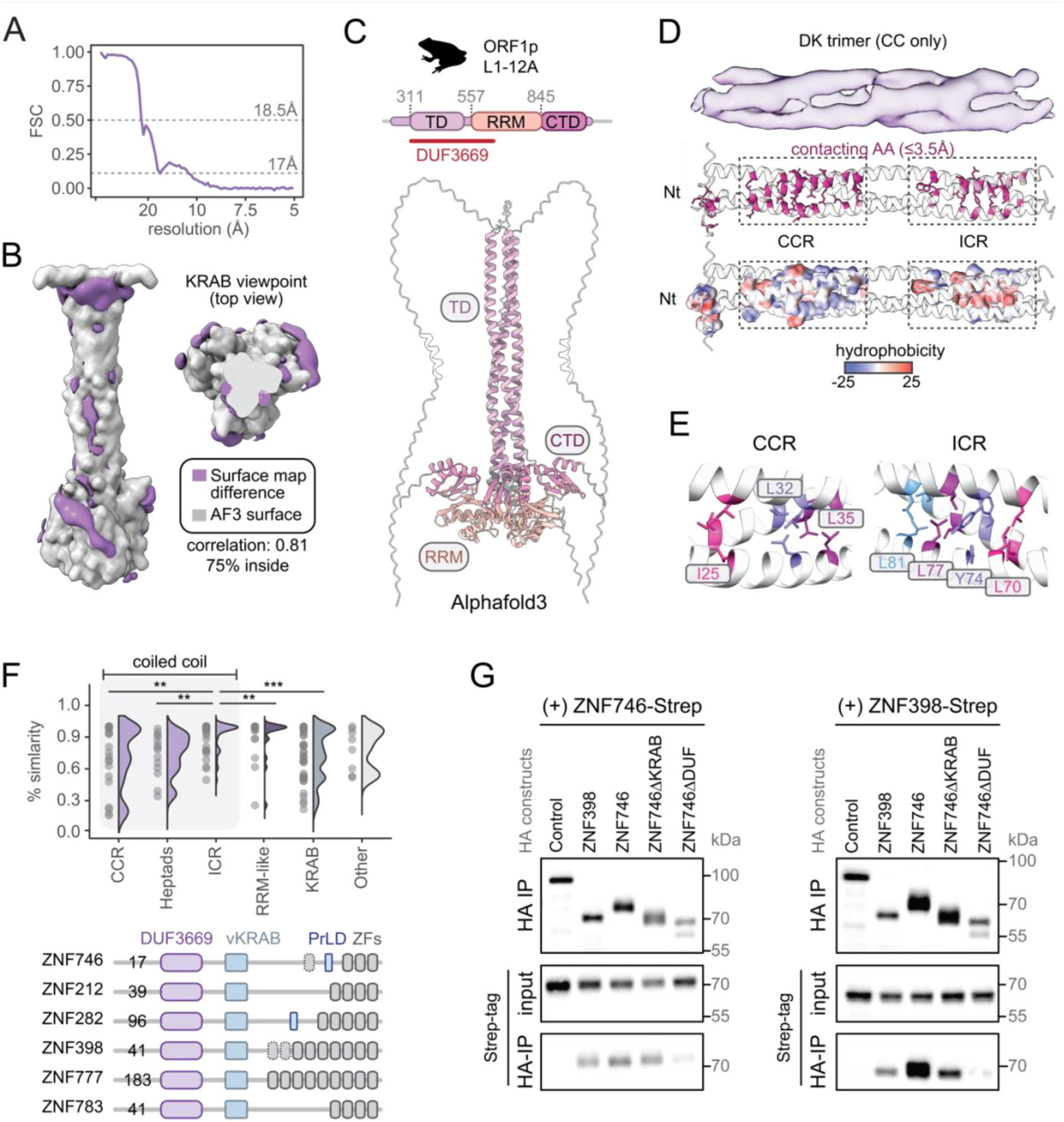
Structural and functional characterization of the DK fragment and its interactions. A. Fourier shell correlation for the DK fragment, related to Fig. 2C. B. Differences (purple) between the AlphaFold3 surface model and the Cryo-EM density map. C. AlphaFold3-Multimer prediction of L1-12A_XT as described in fig. S3E. D. Cryo-EM density map (top) and AlphaFold3-Multimer simulation (two bottom models) of the trimeric coiledcoil structure of the DK fragment. The two AlphaFold3 models illustrate contacting amino acids (≤3.5 Å) colored in pink (middle) or based on their hydrophobicity (Fauchère-Pliska scale, bottom). E. Close-up of amino acid contacts within the CCR and ICR at distances under 3.5 Å, mutated into glutamine. The 5th amino acid mutated in the ICR is highlighted in fig. S4E. F. Percentage of amino acid similarity (top) across the indicated domains of amniote DKZFPs. Each dot represents the mean pairwise similarity at each amino acid position within the indicated domain (Wilcoxon rank sum test). Schematic representation of the six human DKZFPs (bottom), with dotted grey boxes indicating degenerated C2H2 ZF and numbers, the size of the N-terminal region. G. Representative western blot analysis of co-immunoprecipitation assays using HA- and Strep-tagged constructs, as indicated. The assays were performed in HEK293T cells. Related to Fig. 2G.

**Figure S6.**
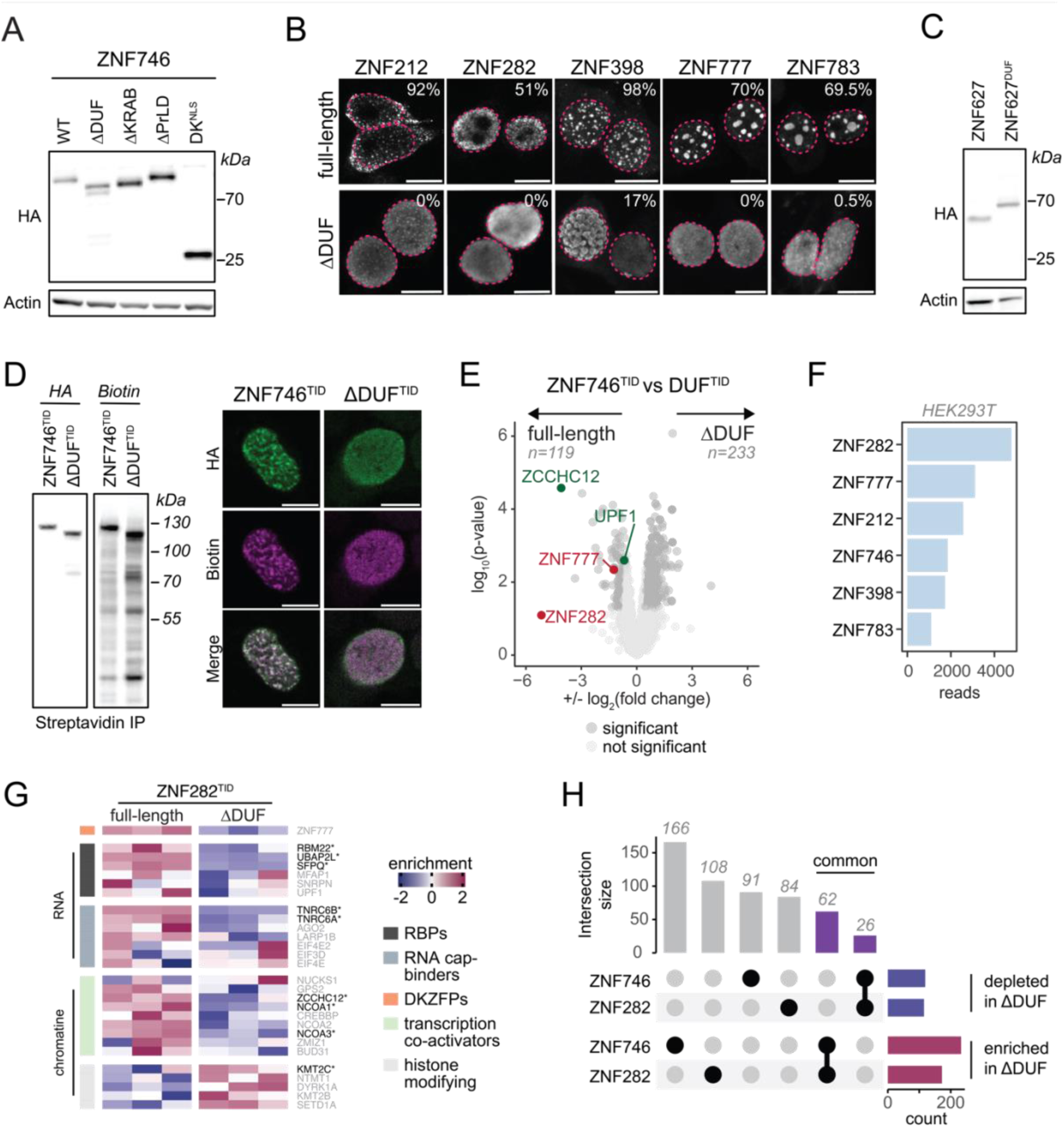
Condensate features vary among DKZFPs. A. Western blot analysis of ZNF746 mutants using anti-HA antibody. B. Representative immunofluorescence images of other HA-tagged DKZFPs and their ΔDUF mutants. Percentages indicate the proportion of cells with condensates. The pink dotted line delineates the nucleus as assessed by DAPI staining. n indicates the number of HEK293T cells counted. Scale bar : 10μm. C. Western blot analysis of ZNF627 and ZNF627^DUF^ using anti-HA antibody. D. Western blot analysis (left) and representative immunofluorescence images (right) of HA-tagged ZNF746^TID^ and ZNF746ΔDUF^TID^ probed with anti-HA and anti-streptavidin antibodies. E. Volcano plot of proteins significantly enriched or depleted in the ZNF746^TID^ versus ZNF746ΔDUF^TID^ comparison. F. Expression levels of DKZFPs in wild-type HEK293T cells. G. Heatmap of proteins associated with ZNF746, identified in the ZNF282 turbo-ID experiment (related to Fig. 3G). Proteins shown in grey were not significant in the ZNF282^TID^ versus ZNF282ΔDUF^TID^ comparison. H. Upset plot depicting the overlap of proteins significantly enriched or depleted for ZNF746 and ZNF282.

**Figure S7.**
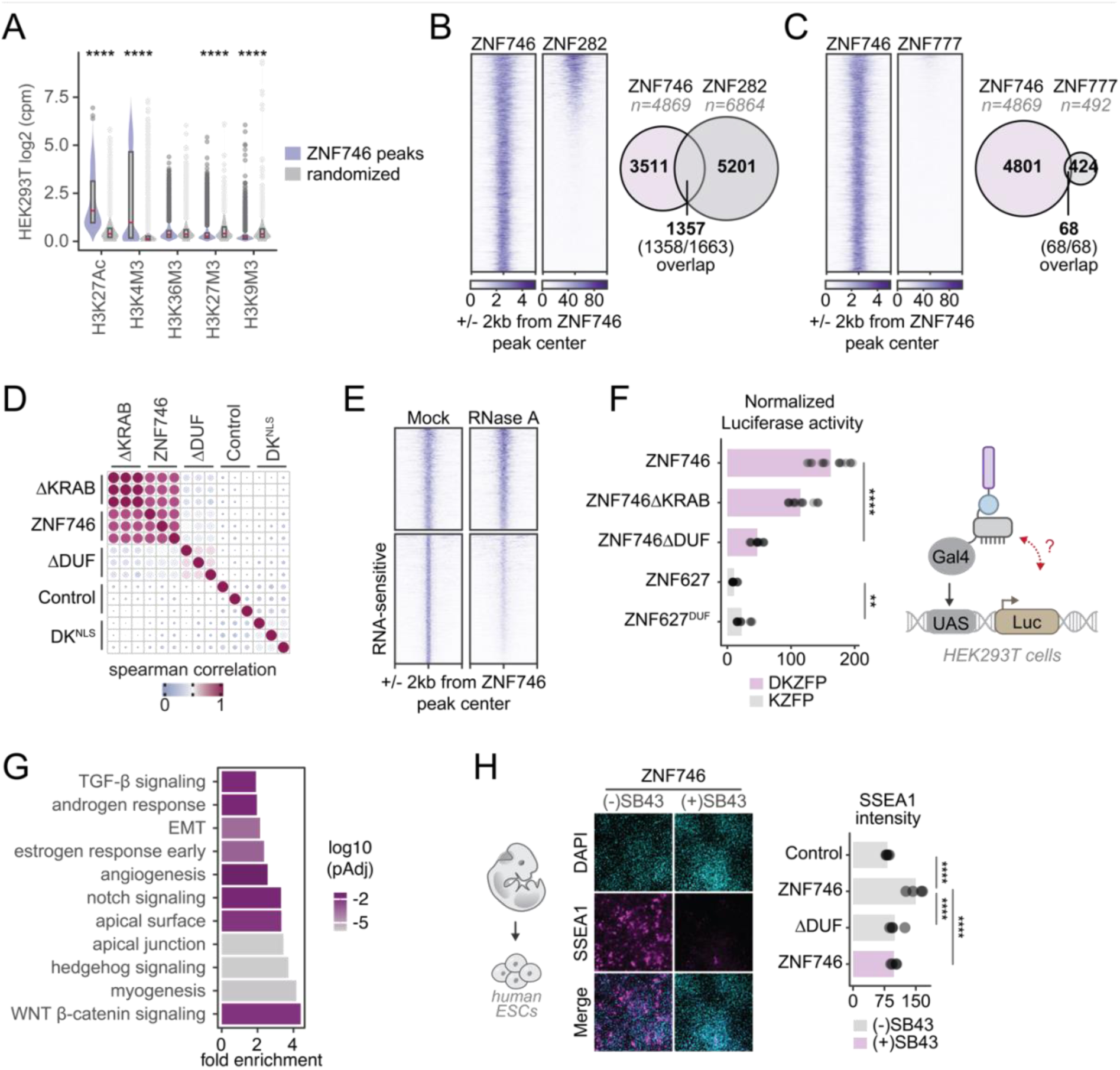
ZNF746 relies on the DUF3669 for genomic affinity and activity. A. Box plot of indicated epigenetic mark enrichment over ZNF746 binding sites or randomized peaks in HEK293T cells (adjusted p-values from multiple Wilcoxon rank sum test). B. (Left) Heatmap of ZNF746 Cut&Tag and ZNF282 ChIP-seq signals over ZNF746 peaks. (Right) Euler plot depicting the overlap of peaks for ZNF746 and ZNF282 (not significant by permutation test). C. Same as B, but with ZNF777. D. Dot plot of the Spearman correlation of Cut&Tag signals in HEK293T cells across three replicates. E. Heatmap of ZNF746 Cut&Tag signals over ZNF746 peaks, with or without RNase A treatment. F. (Left) Schematic of the luciferase assay. (Right) Bar plot of luciferase activity in HEK293T cells (adjusted p-values from multiple t-tests). G. Enrichment analysis of hallmark gene sets using the Molecular Signatures Database for genes near ZNF746 peaks, performed with GREAT (hypergeometric fold change >2, hypergeometric adjusted p-value <0.05). H. Representative images of SSEA-1 staining after 7 days of ZNF746 overexpression, with or without TGF-β signaling inhibition (SB-43, 10 μM) (adjusted p-values from multiple t-tests).

